# Robust and Accurate Doublet Detection of Single-Cell Sequencing Data via Maximizing Area Under Precision-Recall Curve

**DOI:** 10.1101/2023.10.30.564840

**Authors:** Yanshuo Chen, Xidong Wu, Ke Ni, Haoran Hu, Molin Yue, Wei Chen, Heng Huang

## Abstract

Single-cell sequencing has revolutionized our understanding of cellular heterogeneity by offering detailed profiles of individual cells within diverse specimens. However, due to the limitations of sequencing technology, two or more cells may be captured in the same droplet and share the same barcode. These incidents, termed doublets or multiplets, can lead to artifacts in single-cell data analysis. While explicit experimental design can mitigate these issues with the help of auxiliary cell markers, computationally annotating doublets has a broad impact on analyzing the existing public single-cell data and reduces potential experimental costs. Considering that doublets form only a minor fraction of the total dataset, we argue that current doublet detection methods, primarily focused on optimizing classification accuracy, might be inefficient in performing well on the inherently imbalanced data in the area under the precision-recall curve (AUPRC) metric. To address this, we introduce RADO (Robust and Accurate DOublet detection) - an algorithm designed to annotate doublets by maximizing the AUPRC, effectively tackling the imbalance challenge. Benchmarked on 18 public datasets, RADO outperforms other methods in terms of doublet score and achieves similar performance to the current best methods in doublet calling. Furthermore, beyond its application in single-cell RNA-seq data, we demonstrate RADO’s adaptability to single-cell assays for transposase-accessible chromatin sequencing (scATAC-seq) data, where it outperforms other scATAC-seq doublet detection methods. RADO’s open-source implementation is available at: https://github.com/poseidonchan/RADO.

## 1 Introduction

The advent of high-throughput single-cell RNA sequencing (scRNA-seq) has propelled an unprecedented exploration into complex biological systems, unveiling intricate cell heterogeneities and facilitating rare cell-type identifications [1,2,3]. Despite the groundbreaking insights it offers, the reliability of scRNA-seq is constrained by the presence of doublets or multiplets, a technical artifact where two or more cells are encapsulated in a single droplet during library construction [4]. Generally, there are two types of doublets: heterotypic and homotypic [5]. Heterotypic doublets are particularly concerning because they are formed from transcriptionally distinct cells, which can create an artificial new cell type in the dataset. Therefore, doublets may lead to incorrect conclusions in the analysis.

In practice, the best way to mitigate the adverse effects of doublets has been a combination of experimental and computational approaches. Experimental methods [6,7], including sample multiplexing techniques that involve nuclear or cellular surface barcoding, have proven effective in identifying a significant fraction of doublets, thereby enhancing the reliability of downstream analyses. On the other hand, computational methods [8] leverage the principle that if samples are collected from different individuals and they form a doublet, the single nucleotide polymorphisms will present in the same droplet’s sequencing reads. However, these strategies also have limitations, particularly in discerning doublets from cells with identical sample indices or for samples from the same individual, respectively. Furthermore, they are incapable of retroactively filtering out doublets from existing datasets.

In response to these challenges, a variety of computational approaches have been developed to address the issues of doublet detection in scRNA-seq data [9]. Different from the earlier mentioned computational methods, these methods predict whether a droplet is a doublet or not based on scRNA-seq expression data. Proven by previous research, artificially simulated doublets—created by combining two droplets—are quite similar to the real doublets [5]. Consequently, almost all of them adopt the same strategy to simulate artificial doublets and train a classifier on simulated data which is subsequently applied to the real data to predict the presence of doublets. In general, these methods evolve from a basic K Nearest Neighbors (KNN) classifier (e.g., DoubletFinder [5], Scrublet [10]) to more advanced machine learning frameworks like scDblFinder [11] and vaeda [12], which employ the iterative updating and the positive-unlabelled learning strategy, respectively. Importantly, advanced frameworks still consider the predictions from the basic KNN classifier as a crucial feature to boost the prediction performance.

The Area Under the Curve (AUC) encompasses both the area under the receiver operating characteristic curve (AUROC) and the area under precision-recall curves (AUPRC), serving as a widely utilized metric for assessing binary imbalanced classification tasks. Given that doublets typically form a small fraction of a single-cell dataset, AUC has been used to evaluate the performance of these models in previous studies. Nevertheless, when dealing with datasets where the number of negative examples greatly outweighs the positive ones, AUROC may not be the most suitable choice [9,13]. In such scenarios, AUPRC is a more effective choice because it does not depend on true negatives. Yet, current methods predominantly employ cross-entropy as the loss function to optimize parameters. This approach might be suboptimal since cross-entropy is designed as a surrogate function for accuracy, emphasizing the alignment between the choice of loss function and evaluation metric. Therefore, an algorithm that minimizes the cross-entropy loss does not always maximize AUPRC.

In order to tackle the issues arising from data imbalance and the lack of alignment between evaluation metrics and optimization criteria, it appears essential to undergo a paradigm shift. This shift involves prioritizing the direct optimization of the AUPRC metric rather than focusing solely on improving classification accuracy. Recent research [14] has studied AUPRC maximization by using a finite-sum average precision (AP) setting and replacing the ranking function in the AP function with a surrogate loss. This approach maintained biased estimations of the surrogate ranking functions for each positive data point and introduced an algorithm that optimizes AUPRC with guaranteed convergence. Building on this, other studies [15] have developed both adaptive and non-adaptive methods. These methods innovatively updated the biased estimations for each data point and applied momentum averages to both the outer and inner estimators, tracking individual ranking scores. Further advancements in the field [16,17,18] have introduced advanced algorithms that leverage techniques like parallel speed-up and variance-reduction to improve convergence rates. These approaches hold the potential to boost the efficacy of doublet detection tools, facilitating a more accurate annotation of doublets in imbalanced datasets.

In this light, we propose a novel computational algorithm, RADO (Robust and Accurate DOublets detection), based on components analysis and AUPRC maximization, to detect doublets in single-cell sequencing data. It showcases remarkable performance across various real datasets. Our main contributions can be summarized as follows:

- We have developed a computational approach to address the challenges of doublet detection in single-cell sequencing data by maximizing the AUPRC. This effectively tackles data imbalance and enhances model robustness, especially when the simulated data ratio varies and the positive sample ratio is extremely low.
- Through benchmarking on 18 public datasets and using traditionally annotated doublets as the gold standard for evaluation, we demonstrate RADO’s superior performance in terms of doublet score and doublet calling. Using a rigorous statistical test, we show that RADO surpasses all existing methods in the AUPRC metric and achieves performance comparable to the best current methods in the F1 score metric.
- Utilizing 2 public DOGMA-seq [19] datasets, we demonstrate the adaptability of RADO in detecting doublets in single-cell assays for transposase-accessible chromatin sequencing (scATAC-seq) data. Our results suggest that RADO outperforms existing scATAC-seq doublet detection methods when applied to the ATAC modality of the DOGMA-seq datasets.
- In this work, we highlight a promising direction in the field of doublet detection. By maximizing the AUPRC, a simple logistic regression model can outperform state-of-the-art machine learning frameworks, implying significant potential for further performance improvements.

## 2 Methods

### 2.1 The RADO framework

As shown in Fig. 1, the entire pipeline processes raw single-cell data to predict whether each droplet is a doublet or not. The first step involves simulation due to the lack of ground-truth labels for a specific dataset. After the simulation, we compute the KNN doublet score for each droplet. That is, a droplet is likely a doublet if most of its neighbors are doublets. With the KNN doublet score and features extracted from principal component analysis (PCA), we train a logistic regression classifier with AUPRC loss. Then we do it repeatedly to make predictions on the whole dataset in a cross-validation way. We will describe each step in detail in the following part.

**Fig. 1.**
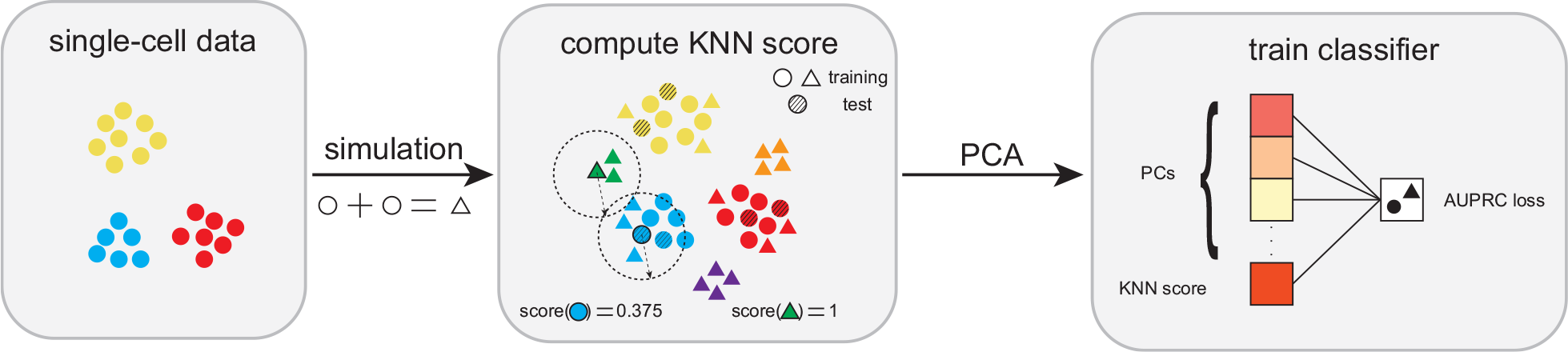
RADO starts with single-cell data, and then simulates doublets by averaging two random droplets. Subsequently, the KNN score is computed and integrated with the top 10 principal components to form the input features. A logistic regression classifier is then trained using the AUPRC loss. The whole dataset’s doublet annotation is finished in a cross-validation way by splitting data into many folds and making training and prediction iteratively.

#### Doublet simulation

Similar to other bioinformatic problems where true labels are usually hard to obtain [20], doublets are simulated by randomly pairing two droplets and then taking their average. In the default setting, the number of simulated doublets equals the number of unlabelled droplets to balance the training data. In the robustness study, the number of simulated doublets varies to test the model’s robustness under the setting of imbalanced training data.

#### Doublet scoring by K nearest neighbors classifier

Inspired by previous works [11,12], we also compute the prediction of the KNN classifier first as an important feature to boost the overall performance. Given the simulated doublets and original droplets as input, we first normalize the library size of all droplets and then log-transform the data to compute principal components (PCs). For single-cell RNA-seq data, we retain the first 30 PCs, while for single-cell ATAC data, we use the first 50 PCs. This is because scATAC-seq data typically has more features than scRNA-seq data, and we use more PCs to retain sufficient information. With the PCs, we compute the nearest neighborhood graph using Euclidean distance adaptively. That is, for a simulated dataset with *n* samples, *k* is defined as 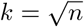. This adaptive KNN computation is proven effective in vaeda [12]. Then, for each droplet, we count the number of simulated doublets (*d*) in its neighborhood to compute the KNN doublet score: 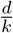. This KNN doublet score intuitively shows the probability of a droplet being a doublet by observing the simulated doublet proportion in its neighborhood. Nevertheless, other dimensionality reduction tools such as auto-encoders could also be effective for this problem [12,21,22].

#### Feature extraction

For single-cell RNA-seq data, we use all the features to compute the PCA. The processing procedure sequentially includes: library size normalization, log-transformation, and PCA. According to previous studies where they have shown that a small number of PCs/latent embedding is good enough for the model [11,12], we only use the first 10 PCs as the input for the classifier. Of note, in our practice, we find that using all raw features leads to better classification performance than that of using highly variable genes only. For single-cell ATAC-seq data, due to the high dimensionality, we use scanpy [23] to select the highly variable peaks (`scanpy.pp.highly_variable_genes(flavor=“seurat_v3”)`) to reduce the running time and memory consumption. We also find that using more ATAC features would lead to better performance. Here, we use the top 25% features to balance the performance and computation complexity.

#### Definition of area under precision-recall curve loss

The AUPRC is a useful performance metric for binary imbalanced data and it is calculated as the area under the PR curve. The definition of AUPRC is given by:

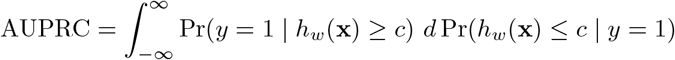

where *h*_*w*_(**x**) is the prediction score function, *w* is the model parameter, (**x**, *y*) is the data point, and *Pr*(*y* = 1|*h*_*w*_(**x**) ≥ *c*) is the precision at the threshold value of c.

In practice, average precision (AP) is used as the estimator of AUPRC to address the issue of the continuous integral in the definition of AUPRC. For a finite set of data set 𝒟 = {(**x**_*j*_, *y*_*j*_), *j* = 1, …, *n*}, it is given by

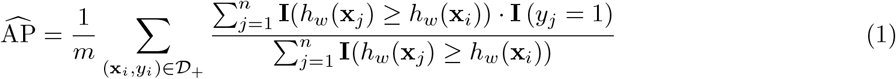

where *m* is the total positive sample number in the dataset, *𝒟*_+_ is the positive data sets, and *h*_*w*_(*x*) is prediction score function with model parameters *w*. Following [14,18], the indicator function **I**(*h*_*w*_(**x**_*j*_) ≥ *h*_*w*_(**x**_*i*_)) is replaced with a squared hinge loss *𝓁* (*w*; **x**_*j*_, **x**_*i*_) = (*max*{*s* − (**h**_*w*_(**x**_*i*_) − **h**_*w*_(**x**_*j*_)), 0})^2^, where *s* is a margin parameter. Finally, we get our optimization function:

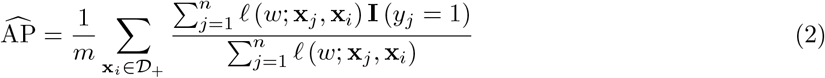

#### Optimization

For convenience, we define the following equation:

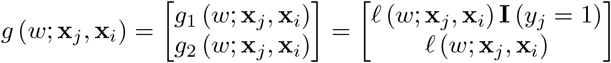

and 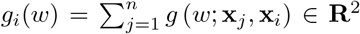 and 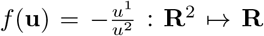 for any **u** = [*u*^1^, *u*^2^]^*⊤*^ ∈ **R**^2^. The final optimization function could be reformulated as the following compositional problem:

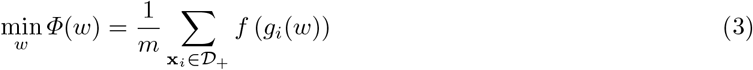

where *m* is the size of positive data sets *𝒟*_+_. Furthermore, the gradient of *Φ*(*w*) is

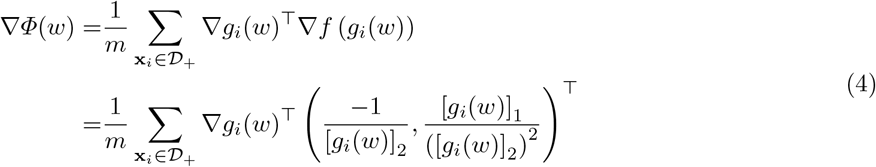

The challenge to optimize the above objective function is we cannot directly use batch sample to get the unbiased gradient estimator because of the two-layer structure in the optimization function and the nonconvex property of the outer-layer function *f*. Following the algorithms in [14], we keep a state [*U*_*t*_] ∈ **R**^*m×*2^ for each positive data point and use recursive momentum to estimate the value of each positive data point during the training. In addition, we define *ĝ*(*w*_*t*_) as below:

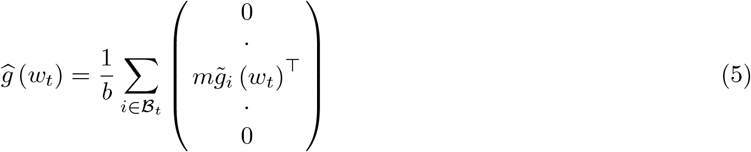

and we have 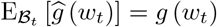. It means both *ĝ*(*w*_*t*_) and 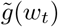 are unbiased estimator of *g*(*w*_*t*_). Then we write the update of *U*_*t*+1_ as 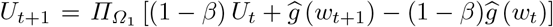, which is equivalent to set 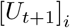 as:

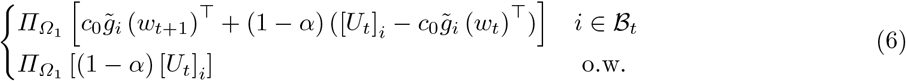

where 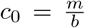. Then we estimate the gradient of the optimization function with the sampled batch size 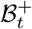, *ℬ*_*t*_ as

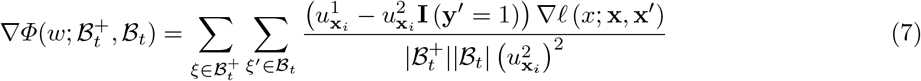

where 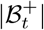 and |*B*_*t*_| denote the sizes of batch 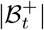 and batch *ℬ*_*t*_, respectively. *ξ* = (**x**, *y*) and *ξ*^*′*^ = (**x**^*′*^, *y*^*′*^). 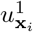 and 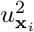 denotes the two values in the state *U*_*t*+1_ for positive data point *ξ* = (**x**_*i*_, *y*_*i*_).

#### Model architecture and training

The classifier is a logistic regression framework implemented in PyTorch [24]. It sequentially comprises one linear layer with bias, one batch norm layer (`torch.nn.BatchNorm1d (1, momentum=0.01, eps=0.001)`) and one sigmoid activation function.

In the training setting, following the previous method [11], we split the whole dataset into 5 folds. Each time, 4 folds of real droplet data along with some simulated doublet data are used as the training set, and we make predictions on the remaining data. After 5 iterations, we can make predictions on the whole dataset. Notably, unlike scDblFinder [11] which iteratively removes doublets in the original datasets, we treat all simulated data as doublets and others as singlets in the training set and validation set. The validation set size is 20% of the whole training set. We use the Adam [25] optimizer with a learning rate 1e-4 to optimize the AUPRC loss of the model. Furthermore, we’ve established early stopping criteria as follows: the training process will terminate only when the AUPRC of the validation set reaches 0.7 or higher, with a 10 epochs patience for early stopping. This heuristic early stopping criterion is proposed after observing that the logistic regression model might underfit certain datasets, given that the AUPRC on the validation set can be significantly below 0.7.

##### Algorithm 1 Maximization of AUPRC

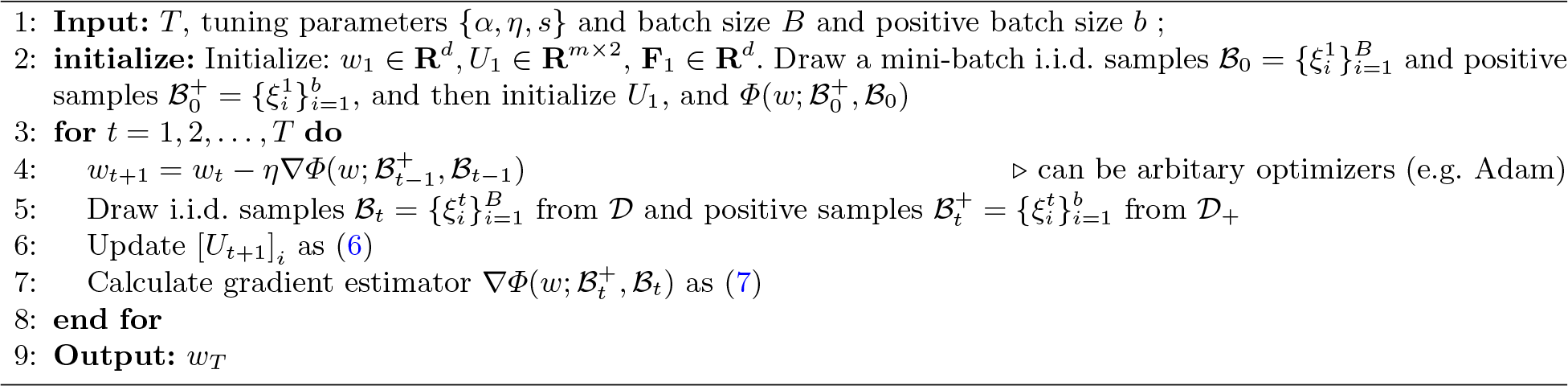

#### Model prediction and doublet calling

After training the model on the training set, the classifier is applied to the remaining data to make predictions. The direct output of the model is the doublet score, which represents the probability of a droplet being a doublet. Unlike the complex doublet thresholding methods used in scDblFinder and vaeda [11,12], we employ a simpler approach by setting a threshold at a probability of 0.5. That is, a droplet with a doublet probability greater than 0.5 is classified as a doublet; otherwise, it is classified as a singlet. After making predictions iteratively, the predictions from each fold are merged to obtain the final prediction on the whole dataset.

#### Hyper-parameter tuning

In the optimization algorithm outlined in Algorithm 1, the model is defined by five tunable hyperparameters: the momentum parameter *α*, learning rate *η*, margin parameter *s*, the number of positive samples *b* in a mini-batch, and the batch size *B*. Initially, we set the margin parameter *s* to 1, aligning with the conventional understanding of hinge loss. The learning rate *η* is set to 1e-4, and the batch size *B* is set to 256, following common practices in scvi-tools [26]. The momentum parameter *α* is set to 0.1 according to the recommendations in the original paper [18]. For simplicity, we make the number of positive samples equal to the number of negative samples in a mini-batch and decide not to tune this parameter during the experiments. Subsequently, we examine the effect of each hyperparameter on the model’s performance. Specifically, we define a range of values for each hyperparameter and evaluate the model’s classification performance across 18 public scRNA-seq datasets for each value. The findings in Table 1 suggest that the model’s performance remains stable across varying values of *α* and *B*. The performance is relatively consistent when the margin parameter is close to 1. A larger or smaller learning rate might harm the training process to lower performance. Finally, the intuitive parameters are kept as the default parameters.

**Table 1.**
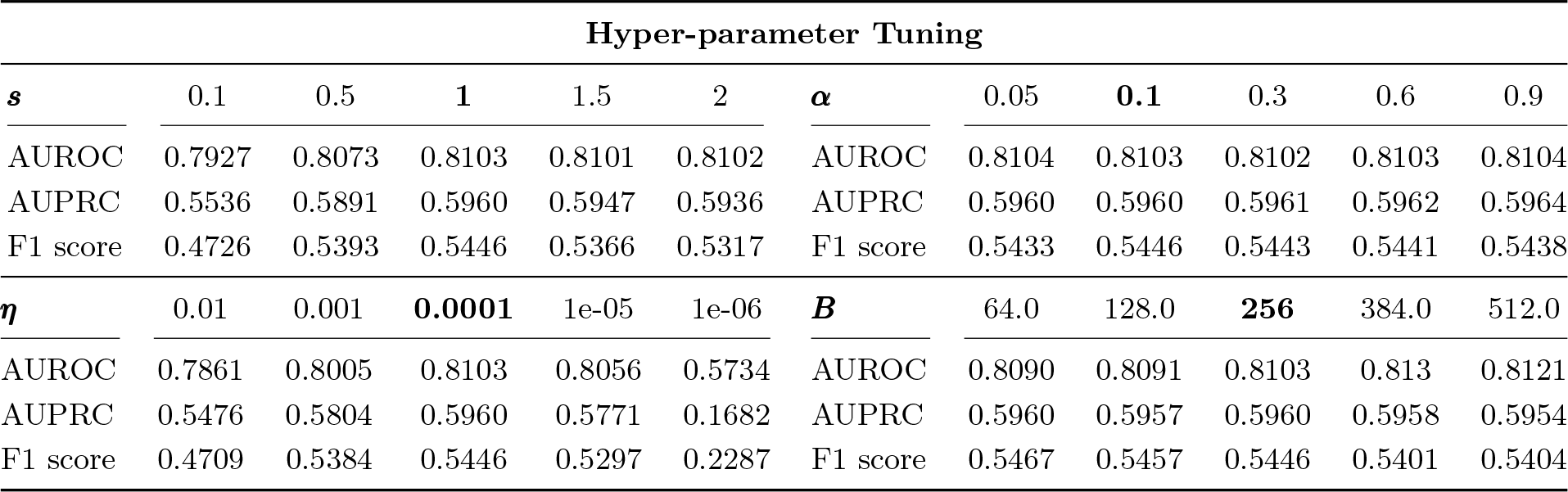
Hyper-parameter tuning experiments show the performance metrics for various hyper-parameter settings. In the experiments, each hyperparameter is individually adjusted while keeping others fixed at their default values. Default values are highlighted in boldface.

### 2.2 Datasets and pre-processing

In this work, the method is evaluated on 18 public real datasets with either experimentally or computationally labeled doublets. Regarding the sequencing data type involved, there are 16 RNA-seq datasets and 2 DOGMA-seq [19,27] datasets. The 16 RNA-seq datasets were collected and processed by a previous benchmarking study [9]. These datasets, namely pbmc-ch, cline-ch [6], mkidney-ch [28], hm-12k, hm-6k [1], pbmc-1A-dm, pbmc-1B-dm, pbmc-1C-dm, pbmc-2ctrl-dm, pbmc-2stim-dm, J293t-dm [8], pdx-MULTI, HMEC-orig-MULTI, HMEC-rep-MULTI, HEK-HMEC-MULTI, and nuc-MULTI [7] are experimentally annotated using sample-hashtag oligonucleotides or genetic variation methods. Specifically, datasets with the suffix “ch” represent the cell hashing technique [6], “MULTI” represents the MULTI-seq technique [7], and “dm” represents for the use of the demuxlet [8] algorithm to annotate doublets with genetic information. Particularly, two datasets, hm-12k, and hm-6k, are annotated by detecting fused human and mouse cell droplets; that is, if one droplet contains sequence reads from both human and mouse, then it is classified as a doublet. However, this method of annotation leads to the ignorance of homotypic doublets, making them two relatively easy datasets. Of note, the doublet ratio of these 16 scRNA-seq datasets ranges from 2.5% (hm-6k) to 37.3% (mkidney-ch). The two DOGMA-seq datasets, which simultaneously measure the expression of ATAC, RNA, and cell surface proteins, are derived from a previous study [27]. These datasets are named as DOGMAseq-DIG-ch and DOGMAseq1-ch, and are annotated via cell hashing. Of note, in this study, only one modality from the DOGMA-seq datasets is utilized in each test. The specific modality used in a particular test is clearly denoted in the name. For instance, “DOGMAseq1-ATAC-ch” indicates the ATAC modality from the DOGMAseq1-ch dataset. Nevertheless, the doublet ratio of these 2 DOGMA-seq datasets is 20.2% and 21.8%, respectively.

### 2.3 Bechmarking and evaluation

For doublet detection methods designed for scRNA-seq data, we include 8 well-known and high-performance methods: scDblFinder [11], vaeda [12], DoubletFinder [5], solo [28], Scrublet [10], and scds [29] which includes hybrid, cxds, and bcds, three different algorithms implemented in the package. For doublet detection applied to scATAC-seq data, we have included three methods: scDblFinder [11], AMULET [30], and ArchR [31]. Our benchmarking procedure aligns with the methodologies employed in prior studies [9,12]. Specifically, we utilize the R package, DoubletCollection [9], which has been developed for benchmarking doublet detection methods, to evaluate scDblFinder, DoubletFinder, and scds on RNA-seq datasets. For Python packages such as vaeda, solo, and Scrublet, we write scripts and execute them using recommended parameters. Regarding the doublet detection for scATAC-seq, we employ the scDblFinder package to conduct experiments for both scDblFinder and AMULET, as AMULET is integrated within this package. Notably, the R implementation of AMULET has been demonstrated to exhibit comparable or superior performance relative to the original Python implementation [11]. We also follow the parameter settings outlined in the scDblFinder tutorial to perform the analysis. For ArchR package, the arrow file is created based on the original fragments file and doublet is detected and filtered with the default parameters.

In this work, the performance of doublet detection is assessed using three metrics: AUROC, AUPRC, and F1 score. Both AUPRC and AUROC are utilized to evaluate the doublet scores due to their relevance in evaluating imbalanced data, a common scenario in the doublet detection task. On the other hand, the F1 score is employed to assess doublet calling, as it is a widely-used metric for evaluating real-world classification performance, providing a balanced measure of precision and recall. We use the evaluation metrics implemented in the Python library scikit-learn [32] throughout the experiments.

## 3 Results

### 3.1 Accurate doublet detection on single-cell RNA-seq data

Doublet detection in scRNA-seq has been extensively studied, and a standard benchmark comprising 16 scRNA-seq datasets has been established by previous benchmarking research [9]. In addition to these 16 well-established datasets, we incorporated 2 DOGMA-seq datasets where only the RNA modality is utilized in the comparison procedure. Hence, we include a total of 18 RNA-seq datasets to comprehensively show the performance of RADO.

The comparison results from Fig. 2 suggest that RADO consistently shows the best average performance across the three evaluation metrics, surpassing the current top method, scDblFinder, by an average of 1 percent. Given the imbalanced datasets, we primarily focus on discussing the AUPRC and F1 score performance rather than AUROC. We find that RADO outperforms all methods on 7 datasets in the AUPRC metric, which is much better than scDblFinder (3), vaeda (2), solo (2), and cxds (3). Meanwhile, DoubletFinder, hybrid, and bcds do not outperform all methods on any dataset. For a more reliable and quantitative comparison, we employ a one-sided paired Wilcoxon test to assess performance differences between pairs of methods. As summarized in Table 2, it is clear that RADO significantly (*p <* 0.05) outperforms all other methods.

**Table 2.** Pairwise comparison of doublet score using the AUPRC criterion. A one-sided paired Wilcoxon test is used to compare the performance of any two methods. For each pair of methods, the direction of comparison (*>* or *<*) indicates that the test is significant (*p <* 0.05) and denotes the relative performance of the method in the row compared to the method in the column. The term “ns” represents that the performance difference between the two methods is not significant (*p* ≥ 0.05).

| Methods | RADO | scDblFinder | vaeda | DoubletFinder | solo | hybrid | Scrublet | bcds | cxds |
| --- | --- | --- | --- | --- | --- | --- | --- | --- | --- |
| RADO | \ | > | > | > | > | > | > | > | > |
| scDblFinder | < | \ | ns | > | > | > | > | > | > |
| vaeda | < | ns | \ | > | ns | > | > | > | > |
| DoubletFinder | < | < | < | \ | ns | > | > | > | > |
| solo | < | < | ns | ns | \ | ns | > | ns | > |
| hybrid | < | < | < | < | ns | \ | > | ns | > |
| Scrublet | < | < | < | < | < | < | \ | ns | ns |
| bcds | < | < | < | < | ns | ns | ns | \ | ns |
| cxds | < | < | < | < | < | < | ns | ns | \ |

**Fig. 2.**
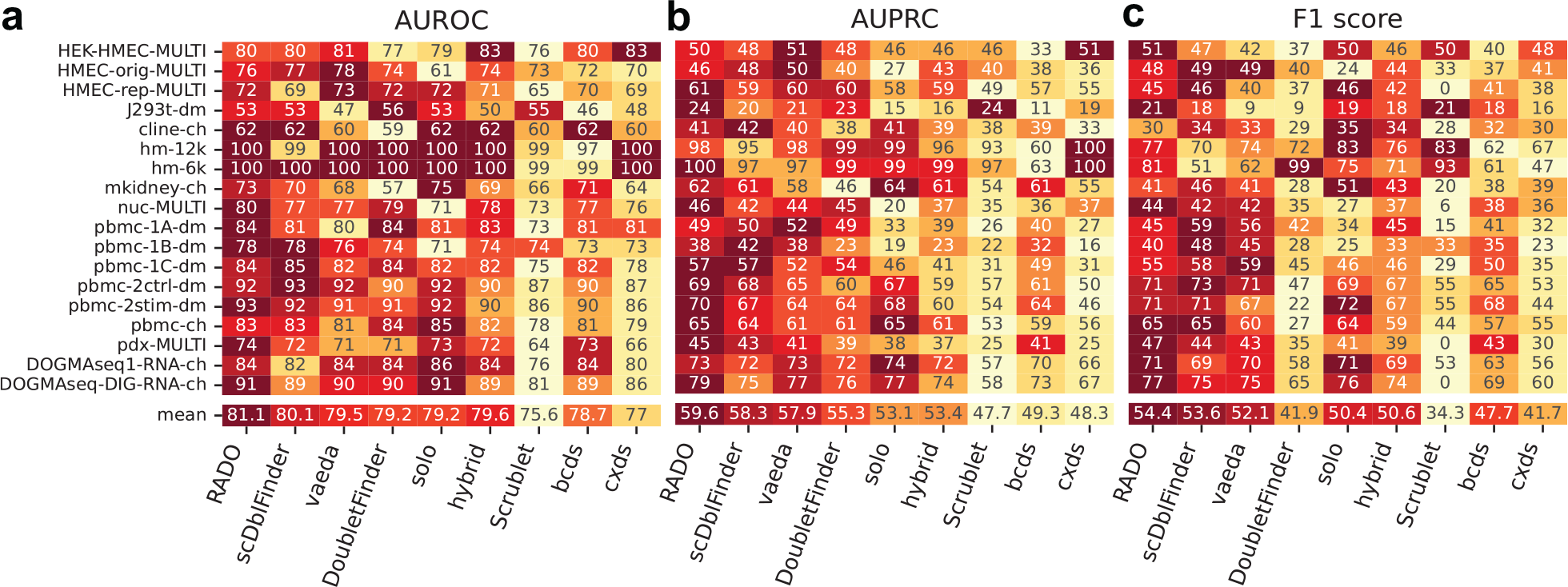
Performance comparison across 18 scRNA-seq datasets. Evaluation metrics encompass AUROC (**a**), AUPRC (**b**), and F1 score (**c**) percentages for each method and dataset. The row labeled “mean” represents the average performance across datasets. In the heatmap, lighter entries represent to lower performance, while darker entries indicate superior performance for each row.

Furthermore, we evaluate doublet calling performance because researchers ultimately need to identify doublets based on doublet scores and filter them out from datasets. In contrast to RADO’s superior performance in the AUPRC metric, even though RADO has the best average F1 score among all methods, scDblFinder and Solo have comparable doublet calling performance, as all three methods outperform all other methods on 5 datasets. Similarly, the quantitative comparison results in Table 3 show that there is no significant difference in doublet calling between RADO and competing methods like scDblFinder and Vaeda. We also observe that DoubletFinder does not perform well in doublet calling, possibly because the doublet calling process heavily relies on the pre-estimated doublet fraction, which varies considerably across the 18 real scRNA-seq datasets.

**Table 3.** Pairwise comparison of doublet calling using the F1 score criterion. Symbols and notation used in this table are the same as in Table 2.

| Methods | RADO | scDblFinder | vaeda | DoubletFinder | solo | hybrid | Scrublet | bcds | cxds |
| --- | --- | --- | --- | --- | --- | --- | --- | --- | --- |
| RADO | \ | ns | ns | > | > | > | > | > | > |
| scDblFinder | ns | \ | > | > | ns | > | > | > | > |
| vaeda | ns | < | \ | > | ns | ns | > | > | > |
| DoubletFinder | < | < | < | \ | < | < | ns | < | ns |
| solo | < | ns | ns | > | \ | ns | > | ns | > |
| hybrid | < | < | ns | > | ns | \ | > | > | > |
| Scrublet | < | < | < | ns | < | < | \ | < | ns |
| bcds | < | < | < | > | ns | < | > | \ | > |
| cxds | < | < | < | ns | < | < | ns | < | \ |

### 3.2 Robust doublet detection at different simulation ratios

While we use simulated data to train the classifier, and although we can create a balanced simulation of positive samples (simulated doublets) and negative samples (unlabeled droplets), we’ve observed that the optimal simulation ratio varies across datasets. Specifically, the best performance in doublet scoring and calling doesn’t always arise from a balanced simulation process. In some cases, better results were achieved when the number of positive samples was fewer than the number of negative samples (Fig. 3a).

**Fig. 3.**
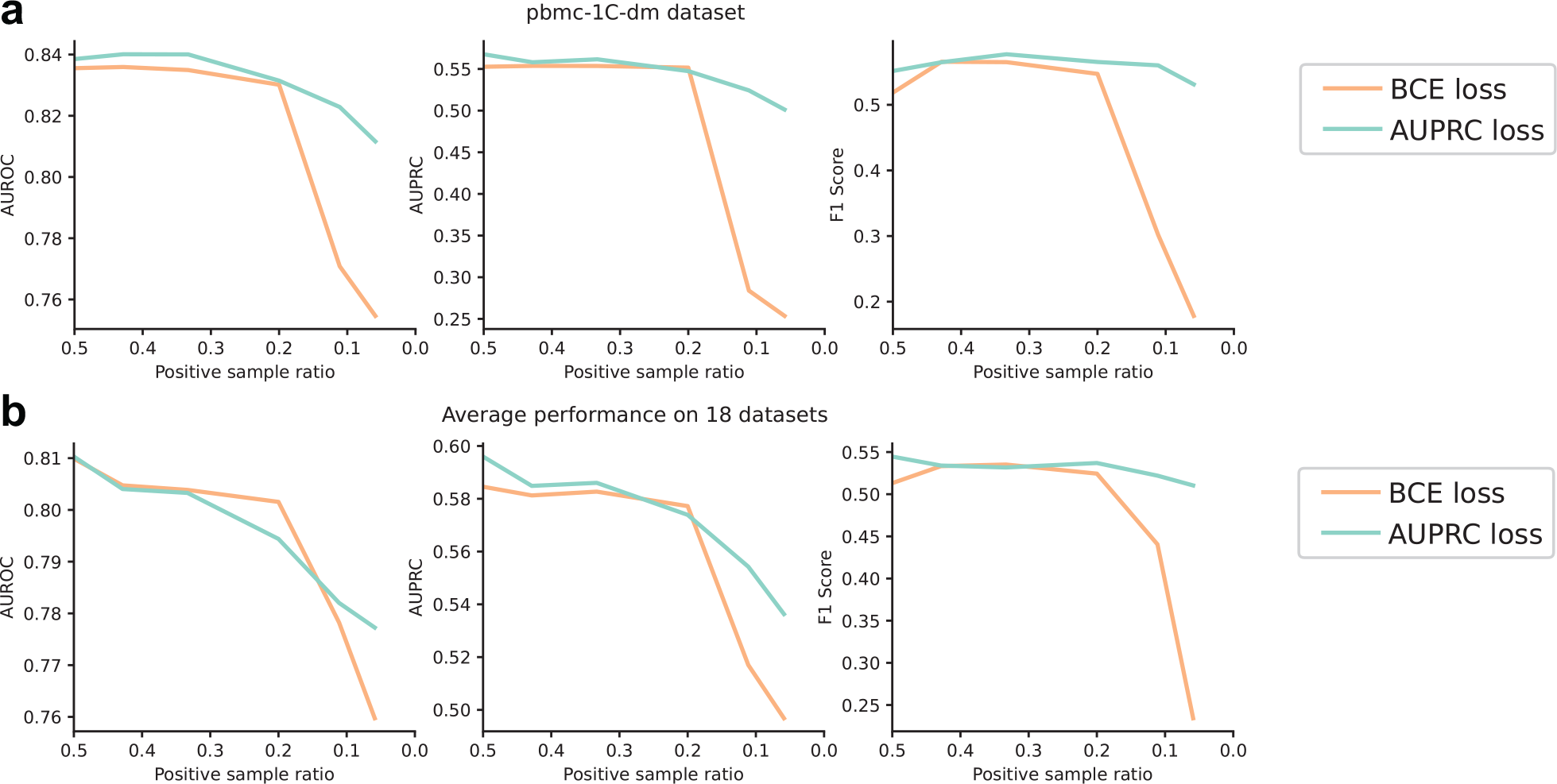
Performance varies at different positive sample ratios. **a** Optimal F1 score performance on the pbmc-1C-dm dataset is achieved with a simulated doublet (positive sample) ratio of less than 0.5. **b** Across 18 scRNA-seq datasets, the average performance shows that training with the AUPRC loss leads to better performance than the BCE loss, especially when the training data is extremely imbalanced.

**Fig. 4.**
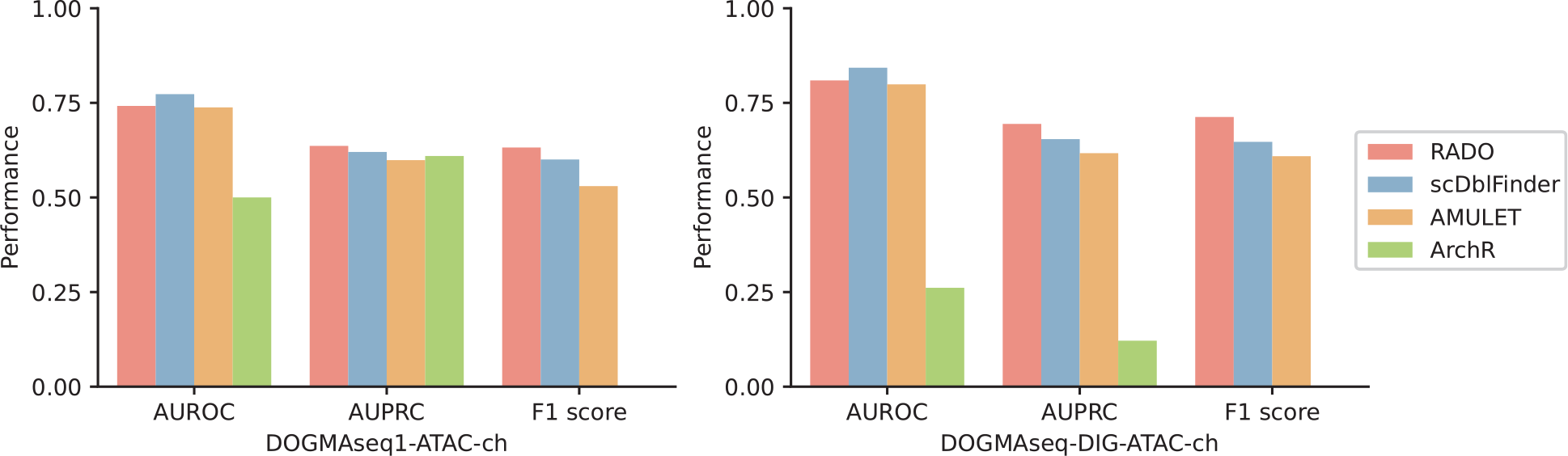
Performance comparison on 2 scATAC-seq datasets. These datasets are subsets from the original DOGMA-seq datasets. In the three metrics, RADO outperforms other methods in terms of AUPRC and F1 score.

This discrepancy might come from our assumption that all unlabeled droplets are singlets. This assumption does not necessarily align with reality, potentially introducing bias in the training process. While the best performance can emerge from an imbalanced simulation process, it’s worth noting that imbalanced data often results in diminished prediction performance [14].

To gain a comprehensive understanding of this phenomenon, we conduct experiments on 18 scRNA-seq datasets and aim to assess the model’s performance at different positive sample ratios and determine whether the AUPRC loss offers any advantage over the traditional binary cross entropy (BCE) loss. The findings, as presented in Fig. 3, indicate that in certain datasets such as pbmc-1C-dm in Fig. 3a, a positive sample ratio less than 0.5 can yield better F1 score performance than balanced training data. However, as a general trend, performance tends to decline as the training data becomes increasingly imbalanced (Fig. 3b). When comparing the two loss functions, the AUPRC loss demonstrates greater robustness in scenarios of extreme imbalance, particularly when the positive sample ratio is below 10%. However, in situations without extreme imbalance, the performance of the two loss functions is quite similar. Interestingly, in Fig. 3b, we also notice that although optimizing AUPRC loss leads to a small AUPRC performance gain compared to the BCE loss, the F1 score gap is really big when the positive sample is rare, indicating the simple thresholding at 0.5 is robust enough when working with the AUPRC loss.

### 3.3 Accurate doublet detection on single-cell ATAC-seq data

Inspired by scDblFinder [11], we realize that the simulation-based doublet detection method could also be extended to other modalities, as long as the simulated doublets closely resemble real doublets. Considering the fact that the ATAC modality can be accurately fused with the RNA modality using state-of-the-art multi-modal integration tools [33,34], which indicates a high similarity between ATAC and RNA data, we propose that our framework, RADO, can also be generalized to scATAC-seq datasets. To demonstrate the accuracy of RADO on scATAC-seq datasets, we benchmarked it against three ATAC-seq doublet detection methods: scDblFinder, AMULET [30] and ArchR [31]. Since scDblFinder, and ArchR are two simulation-based doublet detection methods, the differences lie in feature processing and dimension reduction procedures. Specifically, scDblFinder aggregates peaks at the gene level to make the input identical to the scRNA-seq data for doublet detection, but ArchR utilizes the peak data and uses latent semantic indexing to project data to latent space. In contrast, AMULET is based on the assumption that any given genomic region should be captured at most twice in a diploid cell, and therefore interprets a larger number of loci with more than two reads as indicative of a droplet being a doublet. Though AMULET employs an elegantly simple principle to detect doublets, like ArchR, it also requires the original fragments file as input, which may cause inconvenience for processed matrix-form scATAC-seq data. Unlike other methods, RADO uses the processed peak matrix as input, making doublet detection in scATAC-seq more convenient.

When benchmarking using the ATAC modality from the two DOGMA-seq datasets, we find that RADO consistently outperforms others across the datasets in terms of AUPRC and F1 score. However, scDblFinder shows better performance on the AUROC metric. Comparing this to the performance of doublet detection on the RNA modality of the DOGMA-seq dataset (Fig. 2), we observe that using the RNA modality for doublet detection yields better results than using the ATAC modality. Specifically, RADO’s performance on the RNA modality is 0.73 and 0.79 for these two datasets, respectively, while its performance on the ATAC modality is 0.63 and 0.69, respectively. These results might suggest that the current simulation process is more suitable for the RNA modality. We also observe that ArchR does not perform well on these two datasets, possibly because the sequenced cells are all T cells and the primary type of doublets is homotypic in this dataset, which highly disobeys the assumptions in the ArchR tutorial. Notably, although the simulation-based methods achieve better performance than AMULET, AMULET still maintains its unique advantages in detecting multiplets and distinguishing between heterotypic and homotypic multiplets.

## 4 Discussions

We present RADO, a novel machine learning framework to detect doublet in single-cell sequencing data. Similar to the previous works, RADO synthesizes artificial doublet to train a classifier. Key features distinguishing it from previous methods include (1) highly accurate and robust doublet detection in real single-cell sequencing datasets, and (2) employing the strategy of optimizing the AUPRC loss. In the experiments, we have shown that RADO benefits from the special optimization scheme by maximizing the AUPRC which directly leads to a better performance in the AUPRC evaluation metric compared to the BCE loss. Moreover, optimizing AUPRC loss leads to robustness when the positive sample ratio is extremely low in a classification problem.

Benchmarked on 18 public datasets, we have demonstrated the superior performance of RADO in terms of AUPRC and F1 score. Besides thinking of sample multiplexing techniques [5,35] as the ground truth, we also notice that they have disadvantages in detecting doublet with the same barcode which would produce some false negative results. However, since the simulation-based doublet detection method is not tied to a sample-specific label used in sample multiplexing techniques, we anticipate that RADO might identify some doublets that are false negative in the sample multiplexing methods.

In addition to common scRNA-seq data, we also demonstrate RADO’s adaptability to scATAC-seq data since scATAC-seq and multi-omics data gain more interest these days [19,36]. Although RADO remains superior compared to the state-of-the-art methods, with insights from DOGMA-seq, we observe an intriguing phenomenon: using RNA modality data leads to better performance than using the ATAC modality. While the performance gap can be partially attributed to the feature selection procedure (using only the top 25% of the ATAC data features results in a performance drop), the primary reason might be that simulated ATAC doublets don’t resemble real doublets as closely as simulated RNA doublets do. This observation opens up further discussions on how to simulate and detect doublets in multi-omics datasets.

In conclusion, we have shown that RADO is an accurate and robust doublet detection method for different types of single-cell sequencing data. Considering the fact that RADO has introduced a new paradigm in doublet detection and has the current best performance, we believe that RADO will have a broadly range of users and help enhance the single-cell analysis pipeline. The open-source software can be found at: https://github.com/poseidonchan/RADO.

